# Phosphorylation-State Modulated Binding of HSP70: Structural Insights and Compensatory Protein Engineering

**DOI:** 10.1101/2025.02.17.637997

**Authors:** Mariah Stewart, Chathura Paththamperuma, Colleen McCann, Kelsey Cottingim, Huaqun Zhang, Rian DelVecchio, Ivy Peng, Erica Fennimore, Jay C. Nix, Morcos N Saeed, Kathleen George, Katherine Makaroff, Meagan Colie, Ethan Paulakonis, Michael F. Almeida, Adeleye J Afolayan, Nicholas G Brown, Richard C. Page, Jonathan C Schisler

**Affiliations:** The McAllister Heart Institute, The University of North Carolina at Chapel Hill, Chapel Hill, NC 27599, USA; Department of Pharmacology, The University of North Carolina at Chapel Hill, Chapel Hill, NC 27599, USA; Department of Chemistry and Biochemistry, Miami University, Oxford, OH 45056, USA; Molecular Biology Consortium, Beamline 4.2.2, Advanced Light Source, Lawrence Berkeley National Laboratory, Berkeley, CA 94720, USA; Department of Pediatrics, Children’s Research Institute and Cardiovascular Research Center, Medical College of Wisconsin, Milwaukee, WI 53226, USA; Department of Pathology and Lab Medicine, and Computational Medicine Program, The University of North Carolina at Chapel Hill, Chapel Hill, NC 27599, USA

**Keywords:** HSP70, Phosphorylation, Protein quality control, CHIP (C-terminus of HSC70 interacting protein), Post-translational modifications, Chaperone, Co-chaperone

## Abstract

Protein quality control is crucial for cellular homeostasis, involving the heat shock response, the ubiquitin-proteasome system, and the autophagy-lysosome pathway. Central to these systems are the chaperone homologs heat shock protein 70 (HSP70) and heat shock cognate 70 (HSC70), which manage protein folding and degradation. This study investigated the impact of the C-terminal phosphorylation of HSP70 on its interaction with the co-chaperone CHIP (C-terminus of HSC70 interacting protein), an E3 ligase that ubiquitinates protein substrates for degradation. Using both cell-free and cell-based approaches, including X-ray crystallography, biolayer interferometry, and live cell biocomplementation assays, we demonstrate that phosphorylation at HSP70 T636 reduces CHIP’s binding affinity, shifting the preference toward other co-chaperones like HOP. Structural analysis reveals that phosphorylation disrupts key hydrogen bonds, altering binding dynamics. We engineered a CHIP variant (CHIP-G132N) to restore binding affinity to phosphorylated HSP70. While CHIP-G132N effectively restored binding without additional functional domains, its effectiveness was diminished in full-length phosphomimetic constructs in cell-free and in-cell assays, suggesting that additional interactions may influence binding. Functional assays indicate that phosphorylation of HSP70 affects its stability and degradation, with implications for diseases such as cancer and neurodegeneration. Our findings highlight the complexity of chaperone-co-chaperone interactions and underscore the importance of post-translational modifications in regulating protein quality control mechanisms. By elucidating the molecular details of HSP70 and CHIP interactions, our study provides a foundation for developing therapeutic interventions for diseases characterized by proteostasis imbalance.

## INTRODUCTION

Protein quality control mechanisms are essential for maintaining cellular homeostasis. Components of this system include the heat shock response system (HSR), the ubiquitin-proteasome system (UPS), and the autophagy-lysosome system (1). Together, they balance refolding, degradation, and sequestration, which is essential for proteostasis (2).

Within the HSR, chaperones are the main components that respond to stress and manage substrate outcomes. These components refold damaged or misfolded proteins, fold newly synthesized proteins, and direct proteins to the UPS or lysosome for degradation (2–5). Two major chaperones are the homologs heat shock protein 70 (HSP70) and heat shock cognate 70 (HSC70), differing by only 25%, and are functionally redundant at the protein biochemical level (6). Both bind to hydrophobic regions of non-native proteins and can promote refolding in an ATP-dependent manner (7). HSP(C)70 has three main domains: the nucleotide-binding domain with ATPase activity, the substrate-binding domain, and the C-terminal domain, the lid, with an EEVD tail (6) (**Fig. 1A**). Chaperone outcomes are influenced by co-chaperones that affect the ATPase activity of HSP(C)70. The Carboxyl terminus of the HSC70 interacting protein (CHIP) is an E3 ligase that blocks the ATPase activity of HSP(C)70 and tags substrates with ubiquitin for degradation (8, 9). CHIP has three functional domains: a tetratricopeptide (TPR) domain for binding to chaperones, a coiled-coil domain, and a U-box domain where ubiquitin binds before transfer to a substrate (10) (**Fig. 1A**). The opposing co-chaperone to CHIP is HSP organizing protein (HOP), which also contains multiple TPR domains and increases HSP(C)70’s ATP hydrolysis, enhancing substrate refolding (11). Both co-chaperones interact via the EEVD motif in the C-terminal tail of HSP(C)70, making this domain a significant mechanism in chaperone function and a therapeutic target.

**Figure 1.**
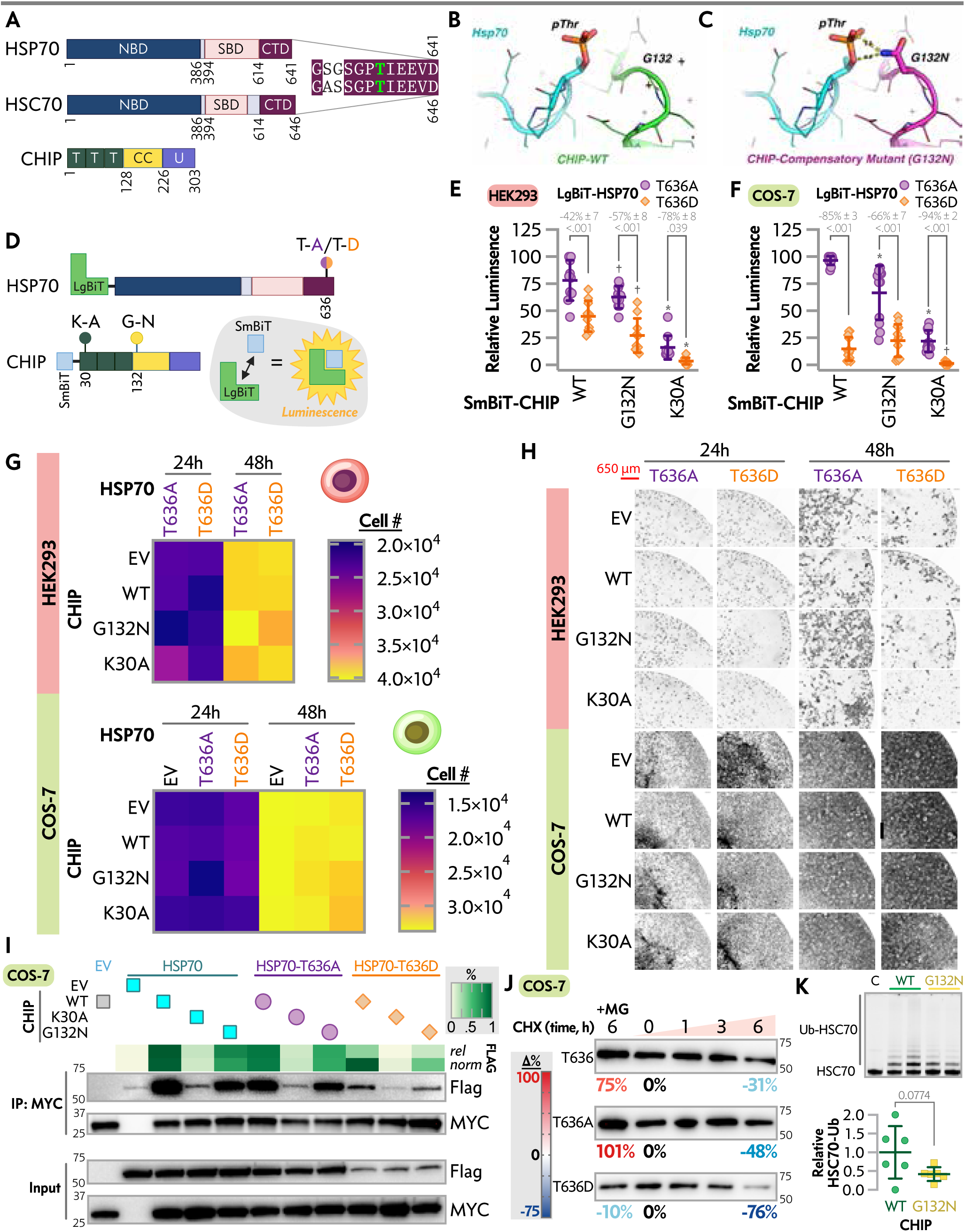
In silico and cell-based characterization of G132N CHIP and HSP70 interactions. *A,* schematics of HSP70, HSC70, and CHIP depicting key domains and amino acid position: Nucleotide-binding domain (NBD), substrate binding domain (SBD), C-terminal domain (CTD), tetratricopeptide repeat (T), coiled-coil (CC), and U-box (U). *B*, *C*, in silico structures of WT CHIP, *B*, and G132N CHIP, *C*, interacting with phosphorylated threonine (pThr-636) of HSP70. D, schematic of NanoBiT protein interaction assay used to assess interactions between LgBiT-HSP70 T636A/D and SmBiT-CHIP variants in HEK-293 and COS-7 cells. *E*, *F*, Percent association of NanoBiT fragments with various CHIP mutants combined with HSP70-T636A or HSP70-T636D in HEK-293 cells, *E,* and COS-7 cells, *F*. Data represented by dot plot and summarized by the mean ± SD from replicates across three independent transfections analyzed via two-way ANOVA: main effects of SmBiT- CHIP, F(2, 54) = 80.93, P < 0.001 and LgBiT-HSP70, F(1, 54) = 63.45, P < 0.001, and an interaction effect between SmBiT-CHIP and LgBiT- HSP70, F(2, 54) = 4.647, P = 0.014. SmBiT-CHIP accounted for 56.08% of the total variation, while the LgBiT-HSP70 accounted for 21.98%. Post-test results within SmBiT-CHIP conditions are included on the plot, * or † indicate P < 0.05 or < 0.001 vs. WT SmBiT. G, cell counts of HEK-293 and COS-7 cells transfected with the indicated HSP70 and CHIP vectors at 24 h and 48 h most post-transfection, represented by heatmap from three independent experiments, results of three-way ANOVA are in Table 1. H, representative images of the conditions in *G*. I, representative immunoblots of MYC (CHIP constructs) and Flag (HSP70 constructs) levels in COS-7 cells detected after immunoprecipitation (IP) of MYC or the cell lysate input. The co-precipitated Flag was quantified via densitometry with relative levels and levels normalized to the Flag input represented by a heatmap. J, representative immunoblots of HSP70 steady-state levels with different amino acids at position 636 following the addition of cycloheximide (CHX) or MG132 with the percent change from time 0 at each 6 h time point. K, representative gel image of cell-free ubiquitination of fluorescently-labeled HSC70 by CHIP-WT or -G132N (upper) and quantification of the relative ubiquitination of HSC70 represented by dot-plot summarized by the mean ± SD from three experiments analyzed via t-test.

Changes in chaperone and co-chaperone interactions affect REDOX control, cell proliferation, and other processes vital for cellular homeostasis and disease prevention. These broad effects have been highlighted in multiple reviews (12–15). Some effects include increased cancer proliferation, neurodegeneration, and heart disease. Proper regulation of this system is essential for homeostasis. For example, CHIP can target tau, a key player in Alzheimer’s disease, but dysregulated CHIP fails to degrade tau, leading to tau aggregates and Alzheimer’s disease development (12).

Additionally, multiple cancer therapies target the HSR, highlighting its role in cancer progression due to improper control (12). One method of regulation is through post-translational modifications (PTMs), which offer potential pharmacological opportunities (15). Understanding these mechanisms and the regulation of CHIP and the 70 family of chaperones is vital for discovering novel techniques to address various diseases. Specifically, determining changes in binding preference between CHIP and HSP(C)70 is a path forward in this endeavor.

Previous studies have shown that phosphorylation of HSP70 affects interactions with co-chaperones like CHIP and HOP (9, 13, 16–18). Phosphorylation changes to the C-terminal tail of HSP70 shift binding affinities away from CHIP to other co-chaperones (13, 16, 18, 19). For example, C-terminal peptides with or without phosphorylation of HSP70 T636 suggest the majority of the phosphorylation-dependent shift is due to a decrease in CHIP binding affinity (16). This prior research highlights the importance of HSP phosphorylation, especially at the C-terminus of HSP(C)70. Still, there is a lack of incorporation of cell-based outcomes and real-time interaction readouts using full-length constructs, and the atomic resolution of these interactions remains elusive. Additionally, most studies have focused exclusively on the tail, not extensively analyzing other interaction points on chaperones, like the lid domain, with co-chaperones like CHIP or HOP (4, 20). Changes in phosphorylation status alter binding and refolding activity, affecting downstream proteins and systems and providing an additional control method for treating diseases mentioned above.

Developing cell-based approaches is crucial for addressing critical gaps identified in the prior research on C-terminal phosphorylation of HSPs and CHIP binding. While previous studies used in vitro assays to mimic phosphorylation and measure binding affinities, cell-based assays offer insights into the dynamic regulation of HSP70 phosphorylation within a cellular context, capturing real-time interaction effects. These assays can validate the physiological relevance of constraining phosphorylation effects on CHIP binding and explore functional consequences on cellular processes like protein degradation and proliferation.

Given the broad and off-target effects of kinase and phosphatase inhibitors, genetic approaches with phosphomimetics and targeted protein engineering to modify protein interactions provide additional means to test mechanistic hypotheses with translational implications. Our study combines cell-based approaches with cell-free studies to understand the impact of C-terminal phosphorylation of the 70 family of chaperones and engineered CHIP on interactions in cells and purified protein systems and the downstream cellular and molecular effects.

## METHODS

### Vectors, peptides, and recombinantly expressed proteins

A complete list of vectors is listed in **Table S1** and deposited at Addgene (pending ID). Plasmids were maintained and propagated in DH5-alpha *E. coli* and purified using the Miniprep plasmid kit (Qiagen) or Quantum Prep® Plasmid Midiprep Kit (Bio-Rad). We used the Q5 Site-Directed Mutagenesis Kit (NEB) or QUIKCHANGE II XL (NC9045762, Agilent Technologies) site-directed mutagenesis kit for editing plasmids and validated via DNA sequencing. Lifetein or Genscript synthesized the peptides used in the study. The unphosphorylated and phosphorylated HSP(C)70 peptide (GPTIEEVD and GP(pT)IEEVD) are termed as EEVD and pEEVD. The amino-terminal biotin-labeled HSP70 tail peptide (biotin-GSGSGPTIEEVD, termed as biotin-EEVD) and the amino-terminal biotin-labeled pT-HSP70 tail peptide (biotin-GSGSGP(pT)IEEVD, termed as biotin-pEEVD) were purchased (Lifetein). The peptides used for fluorescent polarization were N-FITC-Ahx-GSGPTIEEVD (EEVD) or N-FITC-Ahx-GSGP(pT)IEEVD (pEEVD). Recombinant CHIP-TPR constructs were recombinantly expressed using the pHis//2 CHIP-TPR (21–154) plasmid and purified as previously described (21) and additional CHIP proteins were purified as previously described (22).

### Cell Culture

HEK293 and COS-7 cells were acquired from the UNC Lineberger Comprehensive Cancer Center Tissue Culture Facility. Cultures were maintained in Dulbecco’s Modified Eagle Medium with 10% fetal bovine serum at 37 °C and 5% CO_2_. Using the manufacturers’ instructions, X-tremeGENE 9 was used to transfect cells with plasmid DNA except for NanoBiT constructs, in which case we used Lipofectamine 3000 (Thermo Fisher Scientific). Where indicated, twenty-four hours post-transfection, cells were treated with 20 μM MG-132 (Selleck Chemicals) for 6 h or 100 μM cycloheximide (Sigma-Aldrich) for the stated duration.

### NanoBiT Assay

NanoBiT Protein: Protein Interaction System (Promega) was used to measure protein-protein interaction data following the manufacturer’s instructions. As previously described, LgBiT-HSP70 and SmBiT-CHIP construct interactions were used as positive controls (22). Twenty-four hours post-transfection, the cells were collected, replated in 96-well clear bottom white-sided tissue culture plates, and maintained at 37 °C and 5% CO_2_ for 45 min. Nano-Glo Live Cell Reagent (Promega) was added, and luminescence was measured using a CLARIOstar microplate reader over 20 minutes. The total area under the maximum luminescence curve was quantified and used to compare changes in HSP70-CHIP interactions relative to the WT-WT conditions.

### Western Blotting

Samples were collected in a lysis buffer composed of 50 mM Tris-HCl (pH 7.4), 150 mM NaCl, 2 mM EDTA, and 1% Triton X-100, supplemented with 1X Halt protease and phosphatase inhibitor (Thermo Scientific). The samples were then sonicated and combined with Laemmli sample buffer (Bio-Rad). The lysates were resolved on 4-15% TGX Stain-Free gels (Bio-Rad) and transferred to PVDF membranes (Bio-Rad). The membranes were blocked in 5% milk and incubated with primary antibodies (**Table S3**) overnight at 4°C. Following this, HRP-conjugated secondary antibodies (Cell Signaling Technology) were applied. Detection was performed using Lumigen ECL Ultra (Lumigen, tma-100), and the blots were imaged with the ChemiDoc Imaging System (BioRad). Stain-free blots were utilized for normalization, and complete western blot images are provided in **Table S4**.

### Co-Immunoprecipitation

Ezview™ Red Anti-c-Myc Affinity Gel Beads (Sigma-Aldrich) were used for co-immunoprecipitation of myc-tagged proteins. Proteins were eluted via boiling before western blot analysis.

### Cell Counting

HEK293 or COS-7 cells were plated onto 12-well plates. Twelve hours later, the cells were transfected as described previously. Twenty-four hours post-transfection, the cells were replated into clear-bottom, black-walled 96-well plates. Hoechst 33342 nucleic stain was applied to the cells following the manufacturer’s protocol before imaging on an EVOS imager (Thermo Fisher Scientific). Cell counts were determined using the EVOS Analysis Software.

### In-Vitro Ubiquitination Assay

Ubiquitination assays were conducted using recombinant human HSC70, fluorescently tagged on its N-terminus. All assay components except ubiquitin were premixed on ice in a BSA-containing buffer (20 mM HEPES, pH 8.0; 200 mM NaCl; 0.5 mg/mL BSA). Reactions were initiated by adding ubiquitin and were allowed to proceed at room temperature unless specified otherwise. The reactions contained a final concentration of 100 nM Uba1, 1 μM UBE2D2, 200 nM Hsc70, 100 μM Ub, 5 mM MgCl2-ATP, and 1 μM CHIP. The reactions proceeded for 15 minutes before being quenched with SDS loading buffer. The quenched samples were separated by molecular weight via SDS-PAGE in non-reducing conditions using GenScript SurePAGE 4-12% gels. The CHIP-dependent ubiquitination of Hsc70 was visualized and quantified through fluorescent scanning on an Amersham Typhoon 5.

### Fluorescent Polarization Assay

The N-terminally-labelled peptides were loaded into PCR reaction tubes at 2X (40 nM), with the final concentration being 20 nM. Recombinant CHIP constructs were added into the tubes at the indicated concentrations. The reaction was left at room temperature for 30 minutes before loading into black, low-volume, round bottom 384-well plates (Greiner Bio-One) with inter-run triplicates at 18 μl each and subsequently repeated independently two more times. The Clariostar plate reader captured data using an excitation wavelength of 485 nm and an emission wavelength of 530/40 nm, with 100 flashes. The polarization reads were normalized to the dilution buffer. Gain adjustment was made to the tracer HSP70 peptides alone. We used CHIP-WT plus EEVD to calculate Bmax (289) and normalized all polarization data, expressed as % of maximum binding. A nonlinear curve fit for one site/total binding with Bmax constrained to 100 and a constant background of 0 calculated K_d_ (GraphPad Prism v10.4.1).

### Biolayer Interferometry

Bio-layer interferometry measured peptide binding using a BLItz instrument (FortéBio, Sartorius) with streptavidin (SA) sensors. Biotin-EEVD and biotin-pEEVD were solubilized in 20 mM HEPES, 150 mM NaCl, pH 7.0. Affinity measurements were made using the advanced kinetic protocol within the BLItz Pro system software. Biosensors were hydrated for ten minutes in assay buffer comprising 20 mM HEPES, 150 mM NaCl, 5 mM β-mercaptoethanol, pH 7.2. For each affinity measurement, a new, hydrated biosensor was placed on the BLItz tip, and the “Initial Baseline” step was completed by incubation for 30 seconds in 500 μl assay buffer in an opaque black microcentrifuge tube (Argos Technologies). Peptides were loaded during the “Loading Step” by incubating the tip in 4 μl of either peptide for 60 seconds. The “Baseline step” was then conducted by incubation for 30 seconds in 500 μl assay buffer in an opaque black microcentrifuge tube. The “Association step” was performed by incubating the tip in 4 μl of the appropriate TPR protein construct, at protein concentrations of 1 to 25 μM, in assay buffer in an opaque black microcentrifuge tube for 120 seconds. The “dissociation step” was conducted by incubating the tip for 120 seconds in 500 μl assay buffer in an opaque black microcentrifuge tube. Affinities (K_d_) were calculated by the BLItz Pro software as K_d_=k_off_/k_on,_ where k_off_ and k_on_ were determined by individual fits to the “dissociation step” and “association step,” respectively, for each run.

### X-ray Crystallography

Peptides, EEVD and pEEVD, were solubilized in 20 mM HEPES, 150 mM NaCl, pH 7.0. Recombinant CHIP-TPR was mixed with peptides at a 3:1 peptide:protein ratio at a total concentration of 7 mg/ml. Co-crystallization trials were conducted using sitting drop vapor diffusion at room temperature. Crystals of CHIP-TPR/peptides grew in 0.8 μl protein/peptide mixed with 0.8 μl crystallization condition containing 0.2 M lithium sulfate, 0.1 M Tris-HCl, pH 8.5, and 30% (w/v) PEG 4000. Crystals of CHIP-TPR/peptides tail grew in 0.8 μl protein/peptide mixed with 0.8 μl crystallization condition containing 0.2 M lithium sulfate, 0.1 M Tris-HCl, pH 8.5, 30% (w/v) PEG 4000-, and 10-mM zinc chloride.

Crystals were harvested approximately one week after appearance and cryoprotected in LV Cryo Oil (MiTeGen), and frozen in liquid nitrogen before data collection at Lawrence Berkeley National Laboratory, Advanced Light Source beamline 4.2.2. Data were processed in XDS (25). Structures were solved by molecular replacement using PHASER with the structure of CHIP-TPR (PDB ID 4KBQ) (9, 26). Structures were subjected to iterative model building and refined using COOT and PHENIX software packages (27, 28). Geometric and stereochemical validation was performed using MolProbity.

### Molecular Dynamics Simulation

Molecular dynamics simulations were conducted using Gromacs 2020.2 (29). Models for CHIP-TPR/HSP70 tail, CHIP-TPR/pT-HSP70 tail, G132N-CHIP-TPR/HSP70 tail, and G132N-CHIP-TPR/pT-HSP70 tail were generated using the crystal structures solved and reported in this manuscript. The G132N mutation was generated in PyMOL by selecting the m-80° rotamer, a favored backbone-dependent rotamer (30). Initial coordinate files, solvation files, ions, and MD run files were generated using the CHARMM-GUI web interface (31). Simulations utilized a rectangular water box with 10.0 Å padding and sodium and chloride ions at 150 mM concentration. A temperature of 303.15 K was used for the NVT and NPT phases during equilibration. Simulations were conducted using the CHARMM36M force field for 50 ns (32). Trajectories, root mean square fluctuation (RMSF), and hydrogen bond profiles were analyzed using VMD (33). The MD simulation videos can be found at http://hdl.handle.net/2374.MIA/6854

### Statistical Analysis

Normalization and comparisons were performed using GraphPad Prism (v10.4.1) as described in the methods and figure legends.

## RESULTS

### In Silico Engineering of CHIP to Accommodate C-terminal HSP(C)70/90 Tail Phosphorylation

The CHIP-HSP70 interaction is crucial in cellular homeostasis and overall human health and disease (13, 15). In cell-free systems, phosphorylation of the C-terminal tails of HSP70 and HSP90 (**Fig. 1A**) appears to decrease the binding affinity of CHIP to a point that would favor HOP binding (13, 16, 18). Manipulating phosphorylation via kinase and phosphatase manipulation leads to off-target effects that could confound a small molecule approach in examining the mechanism of HSP70/90 tail phosphorylation and co-chaperone interactions (13). We considered a protein engineering approach to modify CHIP to accommodate the phosphorylation event on the C-terminal HSP70/90 tails.

Using a similar in silico approach, as proposed by Muller et al., the phosphorylation of the HSP70/90 C-terminal tail places the phosphorylation sites closer to the CHIP backbone near residues 131 and 132 (16). Seizing on this proximity, we hypothesized that alternative residues at these positions could provide an opportunity to introduce hydrogen bonds with the phosphorylated HSP70/90 C-terminal tails (**Fig. 1B**). Given the role of F131 in interacting with both the HSP70 peptide and α-helical lid subdomain, we focused on residue G132 (37). We identified potential substitutions in silico and found the G132N substitution (**Fig. 1C**) as a candidate for introducing new hydrogen bonds between CHIP and the HSP70 C-terminal tail phosphorylated at T636 (EEVD-pT636).

Using the **m**-80° preferred N side chain rotamer places the side chain of G132N in a position to form two hydrogen bonds (**Fig. 1C**), mediated by the NH2 of the N side chain amide and two of the phosphorylated T636 oxygen atoms (38–40). In contrast, there is no contact between G132N and non-phosphorylated HSP70 (at T636), predicting that G132N will have a similar binding affinity to non-phosphorylated HSP70 as CHIP-WT. If correct, HSP70 could maintain a preference for CHIP-G132N binding following phosphorylation of T636.

### Real-time, In-cell CHIP-HSP70 Interactions and Impact of EEVD Tail Phosphomimetics

Muller et al. used phosphomimetics versions of EEVD tail phosphorylation using A and D substitutions to prevent or mimic phosphorylation, respectively (16). Via pull-down assays using HEK-293 cell lysates, CHIP pulldown levels were highest with the phospho-null, A, for both HSP70 and HSP90 (16). Interestingly, HSP C-terminal tail phospho-null variants slowed cell growth, as measured by electrical impedance (16). Still, the correlation between CHIP binding to HSPs in cell lysates and inhibiting cell growth was unclear and could be due to non-native interactions occurring in cell extracts or CHIP-independent effects of expressing the modified HSP proteins.

To address this, we used a previously established a live cell system to monitor interactions between HSP70 and CHIP through a bio-complementation assay via NanoBiT split (22). We took HSP70 to create a phospho-null (HSP70-T636A) or a phospho-mimetic (HSP70-T636D) version of HSP70 in the NanoBiT system (LgBiT-HSP70-T636A and - T636D, respectively) with CHIP as the interaction partner, including CHIP-G132N, and the K30A substitution known to disrupt the ability of CHIP’s TPR domain to bind EEVD motifs (SmBiT-CHIP-WT, -K30A, - G132N, **Fig. 1D)** (41). Consistent with our hypothesis, HSP70-T636D resulted in a 42% decrease (Δ) in luminescence with CHIP-WT compared to HSP70-T636A (**Fig. 1E**). However, a similar reduction in HSP70-T636D to -T636A was also observed with CHIP-G132N (Δ = 57% decrease, p = 0.2114 vs. Δ with CHIP-WT). As a positive control, CHIP-K30A showed a robust decrease in luminescence with either HSP70-T636A or -T636A. Finally, we observed that CHIP-G132N resulted in lower luminescence with both HSP70-T636A or -T636A when compared to CHIP-WT, suggesting that CHIP-G132N in a HEK-293 live system has overall decreased interactions with HSP70 with similar interaction repercussions regarding the HSP70-T636A/D mimetics. To control for potential confounding effects of endogenous CHIP protein, we validated the real-time system in COS-7 cells and found similar effects of manipulating the T636 residue. Comparing HSP70-T636D to HSP70-T636A resulted in a Δ of −85% and −66% when paired with either CHIP-WT or CHIP-G132N, respectively, with limited luminescence detected when pairing CHIP- K30A with either HSP70 construct (**Fig. 1F**). These data suggest that the phosphorylation of HSP70-T636 impacts transient interactions with CHIP in live cells and that accommodating the phosphorylation via the CHIP-G132N substitution is not sufficient to overcome HSP70-T636 phosphorylation mimicry.

### Effects of HSP70-T636A and HSP70-T636D on Cell Proliferation

In establishing a potential link between the EEVD tail phosphorylation and cell proliferation, previous studies found expressing HSP70-T636A increased HEK-293 cell proliferation, whereas HSP70-T636D had no effect when compared to vector control or HSP70-WT, measured by electrical impedance (16). Given the robust impact of C-terminal tail phosphorylation of HSP70 and CHIP interaction seen in our bio-complementation assays, we next sought to replicate the proliferative potential of HSP70-T636A using live cell counting. Contrary to our expectations, in HEK-293 cells, we did not see any difference in proliferation associated with the HSP70 vector (p = 0.8728), CHIP construct (p = 0.9972), or the interaction between HSP70 and CHIP (p = 0.9997) over time via 3way ANOVA (**Fig. 1G, 1H, Table 1**). For rigor, we performed similar experiments in COS-7 cells, including additional conditions where no additional HSP70 was introduced, and found no effect of HSP70 (p = 0.8691), CHIP (p = 0.9934), or their interaction (p = 0.9959) via 3way ANOVA (**Fig. 1G, 1H, Table 1**). The additional control conditions confirmed that CHIP (p = 0.9993) or HSP70 (p = 0.9969) alone did not change COS-7 proliferation in this cell system.

### Stable Protein-protein Interactions and the Expression and Degradation of HSP70 C-terminal Tail Phosphomimetics

As an orthogonal approach to bio-complementation assays, we performed immunoprecipitations and immunoblot analyses to compare the interaction between CHIP constructs and C-terminal HSP70 tail phosphomimetics. Consistent with the live-cell bio-complementation assays, CHIP-WT co-precipitated similar levels of HSP70-WT and HSP70-T636A, but over 90% less HSP70-T636D (**Fig. 1I**). As expected, CHIP-K30A failed to precipitate appreciable amounts of HSP70 construct above background levels (**Fig. 1I**) consistent with previous studies (42, 43). CHIP-G132N did not override the effect of the HSP70-T636D phosphomimetic (**Fig. 1I**), consistent with the live cell interaction assay results (**Fig. 1F**)

Post-translational modifications, including phosphorylation, can impact protein turnover (44). Previous studies have highlighted that phosphomimetic modifications of HSR components impact the steady-state levels of the proteins (43). We considered an alternative hypothesis that decreased co-precipitation of HSP70-T636D with CHIP could be due to effects on HSP70 steady-state levels (**Fig. 1I**). Using cycloheximide time course experiments in COS-7 cells, we found more robust decreases in HSP70-T636D steady-state levels after inhibiting translation for six hours (a 76% loss) compared to HSP70-WT and HSP70-T636A, likely through the proteasome, given the recovery of protein levels in the presence of the proteasome inhibitor MG-132 (**Fig. 1J**). Replicate experiments were also done in HEK 293 cells and mimicked the decrease of HSP70 T636D steady-state levels seen in the Cos7 cells (**Fig. S1**). This turnover difference is demonstrated in the work done but Muller et al.; however, it was not quantified or further investigated (16). Finally, we used a cell-free ubiquitination system to eliminate the possibility that changes in CHIP’s ubiquitin ligase activity could confound our interpretation of CHIP-G132N interactions with HSP70. We found that ubiquitination of a 70 substrate was not different between CHIP-WT and CHIP-G132N (p = 0.0774), if anything, there was a trend towards decreased ligase activity in CHIP-G132N (**Fig. 1K**).

### Cell-free Binding Kinetics of CHIP and TPR Domains to Phosphorylated and Unmodified EEVD Peptides

The inability of CHIP-G132N to recover binding in the cell-based assays towards a phosphomimetic-modified EEVD tail could be a limitation of aspartate as a phosphorylated threonine mimetic. To test this directly, we generated synthetic peptides to monitor binding via fluorescent polarization (FP) in cell-free assays using recombinant CHIP protein constructs (**Fig. 2A**) as described by Smith et al (45). In this system we found similar decreases in interactions between the phosphorylated EEVD peptide and CHIP-WT or CHIP-G132N (6-10 fold), compared to the unmodified threonine, with overall less binding affinity towards both peptides in comparing CHIP-G132N and CHIP-WT (**Fig. 2B**). As expected, CHIP-K30A had minimal binding with either EEVD peptide, consistent with the TPR domain being the primary contact with the HSP70/90 tail. Interestingly, whereas the TPR domain is necessary and sufficient to drive the EEVD tail interaction, we found that there was minimal change in the presence of the coiled-coil domain; however, moving the Ubox closer to the TPR in the primary structure eliminated EEVD tail peptide interactions with CHIP, suggesting that regions outside the TPR domain of CHIP can influence CHIP-tail interactions (**Fig. 2C**).

**Figure 2.**
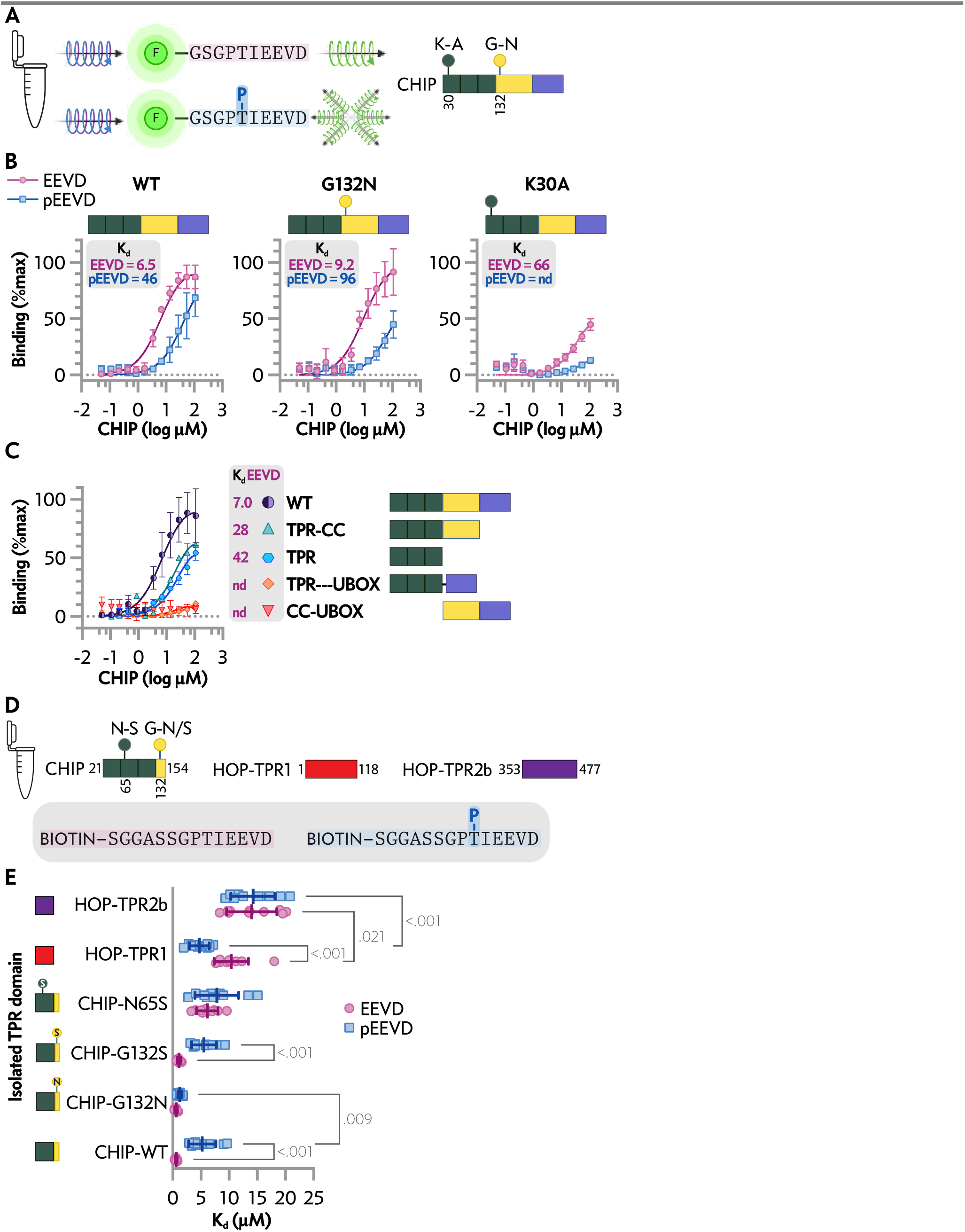
Analysis of EEVD peptide binding to full-length and isolated co-chaperone domains in cell-free conditions. *A, left*, fluorescent polarization of HSP70 peptide tracers (EEVD *upper*, pEEVD *lower*) were used to measure binding affinity to recombinant proteins in a cell-free system, with decreased polarization (lower) indicating less interaction. *Right*, CHIP domains, coded by color: TPR (dark green), coiled-coil (CC, yellow), and the U-box (purple) and locations of key amino acids. *B,* polarization (mP) is represented by the percentage of max binding of EEVD or pEEVD with increasing full-length versions of recombinant CHIP protein: *left*-WT, *center*-G132N, and *right*-K30A. Data are represented by a scatterplot of the mean ± SD from three independent experiments and the resulting curve fit to determine the K_d_. *C,* polarization (mP) is represented by the percentage of max binding of EEVD or pEEVD with increasing CHIP or CHIP domain fragments. Data are represented by a scatterplot of the mean ± SD from three independent experiments and the resulting curve fit to determine the K_d_. *D, upper*, schematic of CHIP and HOP TPR constructs used for biolayer interferometry analysis with biotin-tagged EEVD and pEEVD peptides, *lower*. *E,* binding affinity of isolated TPR domains and EEVD or pEEVD represented by dot plot and summarized by the mean ± SD analyzed via two-way ANOVA: main effects of TRP construct, F(5, 108) = 69.54, P < 0.001 and EEVD peptide, F(1, 108) = 4.584, P = 0.035, and an interaction effect between TPR construct and EEVD peptide, F(5, 108) = 10.95, P < 0.001. TPR construct accounted for 68% of the total variation, while the EEVD peptide accounted for 1%. Post-test results of pairwise comparisons of interest are included in the plot.

FP provides equilibrium binding data by measuring changes in the rotational motion of the fluorescently labeled EEVD when it binds the protein target (**Fig. 2A**), and FP is sensitive to conditions affecting fluorescence, such as pH and temperature. Labeling can sometimes alter binding properties. In contrast, Biolayer Interferometry (BLI) measures the mass of unlabeled molecules binding to a biosensor surface, providing real-time kinetic data and being less affected by environmental factors.

Given the model of CHIP and HOP dictating the outcome of chaperone-bound client substrates, we compared the binding of the HOP TPR domains 1 (residues 1-118), 2b (resides 353-477), and the extended TPR domain of CHIP (residues 21-154) using BLI with EEVD tail peptides on the sensor (**Fig. 2D**). The binding affinity of the CHIP TPR decreased nearly 10-fold towards the phosphorylated EEVD tail of HSP70 (K_d_ = 0.6 vs. 5.2 μM pEEVD vs. EEVD, respectively, p < 0.001 **Fig. 2E**). In contrast, TPR2b of HOP had lower affinity without any difference between phosphorylation status of the EEVD tail (K_d_ = 14.0 vs. 14.3 μM, pEEVD vs. EEVD respectively, p = 0.843, Fig. 2E) whereas TPR1 of HOP did demonstrate binding preference towards the phosphorylated EEVD tail HSP70 (K_d_ = 4.7 vs. 10.4 μM, pEEVD vs. EEVD respectively, p < 0.001 **Fig. 2E**). These data are consistent with previous studies regarding CHIP and HOP binding to the EEVD tails of HSP70 and HSP90, which CHIP being the more sensitive binding partner regarding phosphorylation (16). Finally, we engineered and tested the G132N and additional controls, including G132S (as a site modification control) and N65S, which we previously characterized as a substitution that disrupts CHIP- HSP70 interactions. As expected, CHIP-N65S reduced binding towards both EEVD tails with no preference for phosphorylation status (Fig. 2E). However, CHIP-G132N rescued the binding affinity caused by the phosphorylation of the EEVD tail (K_d_ = 0.6 vs. 1.2 μM, pEEVD vs. EEVD respectively, p = 0.568, **Fig. 2E**) whereas CHIP-G132S exhibited binding affinities similar to CHIP-WT (**Fig. 2E**). Further analysis of the binding constants suggests that the increased binding affinity of CHIP-G132N towards pEEVD may be due to higher association rates (**Fig. S2**).

### Impact of Phosphorylation on HSP70 and Tail Interactions with CHIP TRP Domain

To understand the structural interactions of HSP70 and CHIP at the atomic level, we completed structural studies of the CHIP-TPR domain of CHIP (21–154) with either the phosphorylated or unmodified tail of HSP70. These structures follow our prior structure of the CHIP-TPR in complex with the HSP70 substrate binding domain (SBD), α-helical lid subdomain, and the tail domain, which highlighted residues within the CHIP-TPR that contact only the tail domain, only the α-helical lid subdomain, or both the tail and α-helical lid subdomain (**Fig. 3A**). Although the new structures of CHIP-TPR with HSP70 peptide and pT- HSP70 peptide are globally similar and nearly identical, key differences are observed for T636 and neighboring HSP70 peptide residues. Upon phosphorylation, rotation of the HSP70 peptide eliminates a hydrogen bond between the side chain carboxylate of E638 of HSP70 and the side chain amide of CHIP-TPR Q102 (**Fig. 3C, Fig. S3**). This rotation also causes a minor change to the side chain rotamer for I637 with minor adjustments to the packing of the gamma and delta methyl groups against the hydrophobic surface presented by residues V94 and F98 of CHIP-TPR. Interestingly, the rotation of the phosphorylated HSP70 peptide provides a conformation nearly identical to that of the HSP70 tail within the CHIP-TPR/HSP70 SBD-tail structure (PDB ID 4KBQ). Yet, CHIP-TPR Q102 is rotated away from E638 in our structure with pT- HSP70 (9).

**Figure 3.**
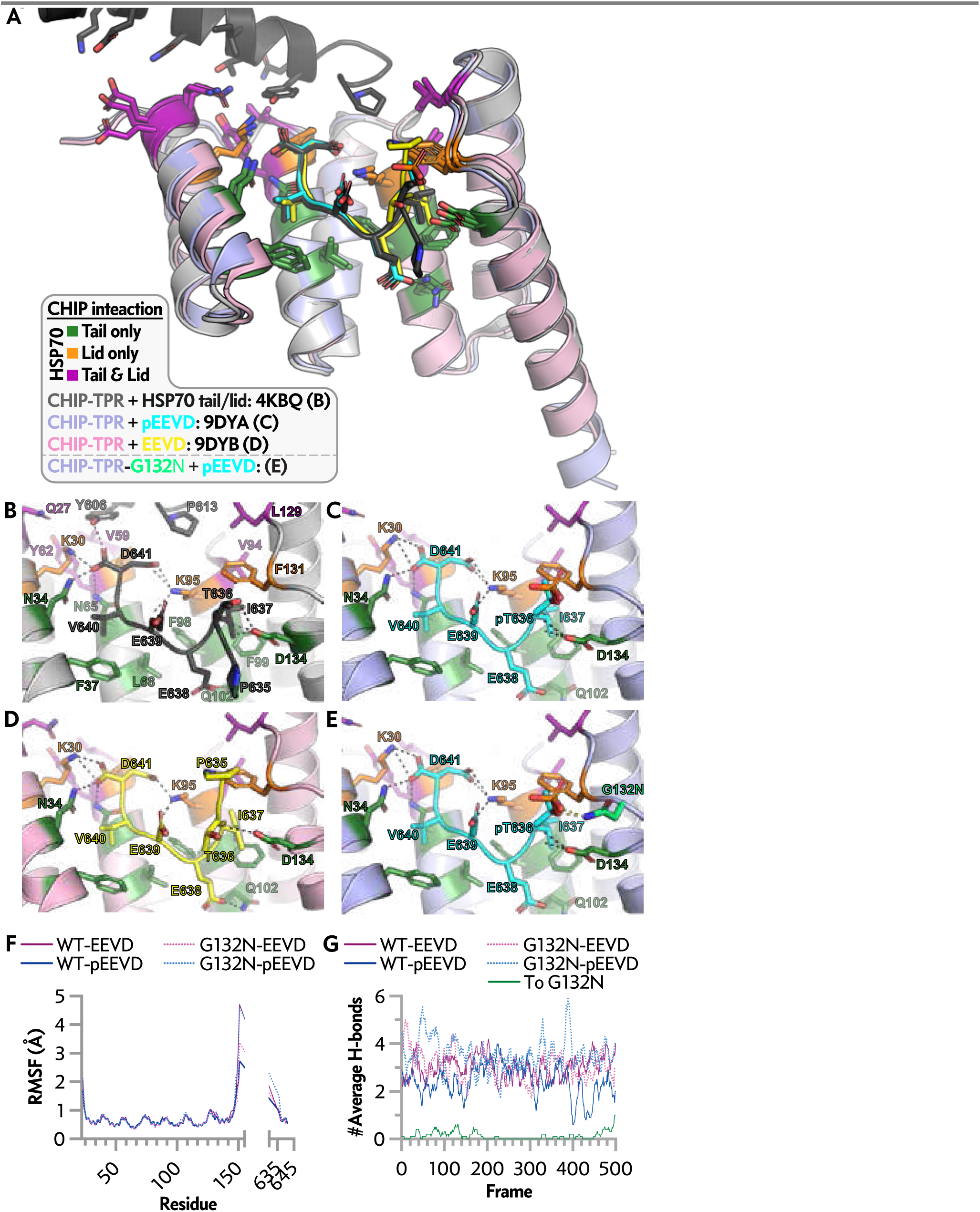
Peptide structures and molecular dynamics simulations show recovery of H-bonds at G132 that accommodate HSP70 tail phosphorylation. *A,* overlay of CHIP-TPR/HSP70 tail peptide structures with the previously solved structure of CHIP-TPR in complex with the helical lid+tail domain of HSP70. PDB IDs and coloring: 4KBQ: CHIP in light grey with HSP70 helical lid+tail domain in dark grey; 9DYA: CHIP in light purple with EEVD in yellow; 9DYB: CHIP in light pink with pEEVD in cyan. Throughout, CHIP-TPR residues interacting with only the HSP70 tail are colored forest green, CHIP-TPR residues interacting with only the HSP70 helical lid domain are colored magenta, CHIP-TPR residues interacting with both the HSP70 helical lid and HSP70 tail are colored orange. *B,* structure of CHIP-TPR in complex with the helical lid+tail domain of HSP70. *C,* structure of CHIP-TPR in complex with pEEVD. *D,* structure of CHIP-TPR in complex with EEVD. *E,* structure of CHIP-TPR with the in silico-modeled G132N mutation in complex with pEEVD with the N132 side chain colored lime green. *F,* comparison of RMSF for C atoms by residue for each trajectory of complexes formed by EEVD (purple) or pEEVD (blue) with CHIP-TPR (solid lines) or CHIP-TPR-G132N (dotted lines). *G,* comparison of intermolecular H-bond profiles for each complex formed by EEVD (purple) or pEEVD (blue) with CHIP-TPR (solid lines) or CHIP-TPR-G132N (dotted lines). Additionally, the contribution of N132 to the total H-bond profile between CHIP-TPR-G132N and pEEVD is plotted (To, green: ten-frame-average number of H-bonds). The MD simulation videos can be found at http://hdl.handle.net/2374.MIA/6854.

In comparison, Q102 is rotated toward and engaged in a hydrogen bond with E638 in the CHIP-TPR/HSP70 SBD-tail structure. Thus, a minor shift of E638 allows for a rotation of Q102 away from the HSP70 peptide, thereby eliminating a hydrogen bond. These minor changes suggest flexibility in the side chain rotameric states that Q102 may adopt, thus making it available to be sampled in solution by dynamic motions. This indicates that structural changes within both the HSP70 peptide and CHIP-TPR may serve as determinants for binding and that minor variations in structure may dictate or allow for the loss or reintroduction of hydrogen bonds that, in part, regulate the affinity of the interaction between CHIP and HSP70. In contrast to the CHIP-TPR/HSP70-SBD structure (**Fig. 3B**) and CHIP-TPR/pT-HSP70 peptide (**Fig. 3C**), the side chain of T636 for the CHIP-TPR/HSP70 peptide (**Fig. 3D**) points away from D134 of the CHIP-TPR, yet a allows for maintaining a hydrogen bond to the T636 backbone, similar to the CHIP-TPR/HSP70-SBD and CHIP-TPR/pT-HSP0 peptide structures. This indicates two key points. First, the position of the HSP70 tail near T636 is different across the structures, indicating the potential flexibility of this region, the potential for this region to adopt more than one conformation, or the presence of crystal contacts forcing one conformation or the other in various crystal packing arrangements. Second, the position of the T636 side chain in the CHIP-TPR HSP70 peptide complex is not available to pThr636 in the CHIP-TPR pT-HSP70 peptide complex because this would place the phosphate group of pT636 in close proximity to D134 of the CHIP-TPR. Consequently, the effects observed near E638 and T636 indicate phosphorylation at T636 removes two hydrogen bonds.

With the structure of CHIP-TPR/pT-HSP70 resolved, we examined the area around pT636 to explore the impact of the G132N substitution. Using an in silico model and applying the most preferred asparagine side chain rotamer for N132, our two desired hydrogen bonds returned (**Fig. 3E**). The NH2 of the asparagine side chain amide makes hydrogen bonds to two of the pT636 oxygens. If this mutation is made in CHIP with non-phosphorylated HSP70, there is no contact between N132 and HSP70, so modeling predicts that G132N should not affect non-phosphorylated HSP70. Due to the position of the N132 side chain, removing the pT636 phosphate group would not result in any steric clashes; therefore, the EEVD tail structure could return to that seen in the CHIP-TPR/HSP70-SBD or CHIP-TPR/HSP70 peptide structures. In fact, these data are consistent with the BLI data, suggesting that in the presence of CHIP-G132N, HSP70 would no longer preferentially shift to other co-chaperones following phosphorylation of T636.

### Molecular Dynamics Simulations: Recovering H-Bonds at G132 Accommodates Phosphorylation

Molecular dynamic simulations were conducted to investigate further the impact of phosphorylated and non-phosphorylated HSP70 binding with CHIP-WT or -G132N, to test our hypothesis that focusing on G132 as a critical residue in phospho-EEVD sensitivity. The average structure may not represent the transient H-bonds that may occur between CHIP-TPR and the peptide. Confirming that the introduced G132N mutation is primarily responsible for the higher affinity in the G132N variant for pT-HSP70 peptides is vital. Accordingly, we analyzed the root mean square fluctuation (RMSF) and intermolecular H-bond profile throughout the simulation (**Fig. 3F, 3G**). The RMSF plot indicates minimal fluctuation of the CHIP-TPR and peptide structures throughout the simulations, except for expected more significant fluctuations at the N- and C-termini and CHIP-TPR and the N-terminal residues of the Hsp70 peptides. A clear pattern of oscillating fluctuations by residue for the CHIP-TPR indicates more stably structured helices linked by comparatively and moderately more flexible interhelical loops. The H-bond profile indicates that CHIP- G132N forms one or more H-bonds in most frames compared to CHIP-WT, suggesting that the molecular mechanism for stronger affinity is a more robust H-bond network. To assess the contribution of N132 to the CHIP-G132N variant, the H-bonds formed by N132 were also plotted (**Fig. 3G**). The results demonstrate that when N132 contributes to creating an H-bond, the total number of H-bonds increases by at least one (**Fig. 3G**, frames 0-200 and 300-500). When N132 does not contribute to forming any H-bond, the total number of H-bonds remains the same as that of the wild-type CHIP-TPR complex with phosphorylated HSP70 peptide. This confirms that the higher affinity of CHIP-TPR(G132N) toward pT-HSP70 is mainly caused by introducing one additional H-bond between the phosphate group and N132.

## DISCUSSION

Our study provides important insights into how phosphorylation affects the binding of HSP70 and its interaction with the co-chaperone CHIP. The findings highlight how post-translational modifications, especially phosphorylation, influence the binding dynamics and functional outcomes of chaperone/co-chaperone interactions. Specifically, phosphorylation of HSP70 at the C-terminal EEVD tail, specifically T636, decreased its binding affinity to CHIP. This is consistent with previous studies showing how phosphorylation can regulate chaperone interactions. Structural analysis revealed that phosphorylation leads to conformational changes that disrupt critical hydrogen bonds, thus reducing CHIP’s binding affinity (**Fig. 3**). This mechanism highlights the importance of phosphorylation as a regulatory switch that can alter chaperone function and cellular proteostasis.

We used a protein engineering approach to study the reduced binding affinity caused by phosphorylation. CHIP-G132N was designed to introduce new hydrogen bonds with the phosphorylated Hsp70 tail. While in silico models predicted successful restoration of binding affinity and validated by BLI experiments using isolated regions of CHIP, experimental data showed that in the context of full-length CHIP, the G132N substitution did not fully compensate for the changes caused by phosphorylation or phosphomimetics both in cell-free systems and in cells (**Fig. 2**). This suggests that other factors or interactions in cells may stabilize the CHIP-HSP70 complex and highlight the complex nature of chaperone-co-chaperone interactions. The coiled-coil domain of CHIP may also play a role in the interaction with HSP70 by influencing CHIP’s folding or dimerization. Further investigation is needed to understand this domain’s role in the HSP70-CHIP interaction.

Functional assays conducted in HEK-293 and COS-7 cells provided insights into the biological relevance of these findings (**Fig. 1**). The phosphorylation-mimetic HSP70-T636D consistently exhibited decreased interaction with CHIP in both cell lines. However, this reduced interaction did not significantly affect cell proliferation as seen in other studies (16, 46), nor did increasing levels of other HSP70 constructs, suggesting that other compensatory mechanisms may be at play to maintain cellular homeostasis. Additionally, the turnover rates of the HSP70 mimetics suggest that phosphorylation may also influence protein stability and degradation pathways (**Fig. 1J**). This underscores the multifaceted role of post-translational modifications in regulating chaperone function that could affect cancer cell proliferation (16). A critical limitation of the cell-based study is the functional differences between an aspartate residue and a phosphorylated tyrosine in mammalian cells. In our cell-free FP and BLI experiments, phosphorylated HSP70 tail peptides were used instead of the cell-based, full-length mimetic (HSP70-T636D). Despite previous work that has extensively used the D mimetic, others have highlighted that the phosphomimic and phosphorylated protein versions behave differently (16, 47–50).

Beyond the interaction between the tail of HSP70 and the TPR domain of CHIP, the flexibility of the HSP70-CHIP complex may also depend on interactions between the lid domain of HSP70 and the TPR domain of CHIP. The chaperone’s ADP/ATP status might mediate these interactions, adding another layer of complexity to regulating this complex.

Our study expands our understanding of how post-translational modifications regulate chaperone function and protein quality control. It emphasizes that while protein engineering approaches like CHIP-G132N offer promising avenues for restoring binding affinity, the complex nature of chaperone-co-chaperone interactions requires further investigation. This includes exploring structural guided design for candidate molecules that bind CHIP-TPR and biophysical analyses of inhibition of these Lid/TPR interactions. The functionality of HSP70 and HSP90 can be regulated by PTMs, collectively called the “chaperone code” (51, 52). A vast number of PTMs have been reported so far for HSP70, which includes phosphorylation. Therefore, the interactions of HSP70 family proteins might be greatly influenced by multiple PTM sites (51). Moreover, HSP70 is not the only component in this system that can be post-translationally modified by phosphorylation. CHIP and its counterpart HOP can also be phosphorylated, altering the interaction between these co-chaperones and chaperones like HSP70 and HSP90 (43, 53, 54). Further investigation of this phosphorylation and its impact on the overall heat shock mechanism could provide insights into how we can pharmacologically control the stress response.

A deeper understanding of these mechanisms will be crucial for developing novel therapeutic interventions to modulate chaperone function in diseases characterized by proteostasis imbalance, such as neurodegenerative disorders and cancer. Future studies should explore the potential of combining protein engineering with small-molecule modulators to fine-tune chaperone interactions. Additionally, investigating other post-translational modifications and their interactions with phosphorylation events could provide a more comprehensive understanding of chaperone regulation.

## ACKNOWLEDGMENTS

The authors acknowledge funding from the National Institutes of Health through R35GM128595 (R.C.P), R35GM128855 (N.G.B.), R01AG066710 (J.C.S.), and R01AG061188 (J.C.S.); and the American Heart Association through 23IPA1048749 (N.G.B.)

## DECLARATION OF GENERATIVE AI AND AI-ASSISTED TECHNOLOGIES IN THE WRITING PROCESS

In preparing this manuscript, we utilized generative AI and AI-assisted technologies to enhance the clarity and conciseness of the text.

Specifically, we employed Grammarly and Microsoft Copilot to provide suggestions and improvements. These tools assisted in refining the language, ensuring grammatical accuracy, and optimizing the overall readability of the document.

