## Supporting Information for "Phosphorylation-State Modulated Binding of HSP70: Structural Insights and Compensatory Protein Engineering"

#### Supporting Figures

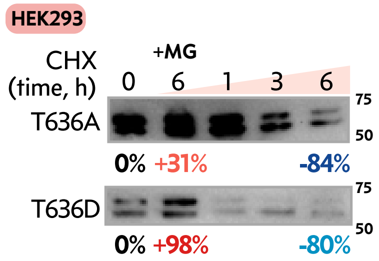


**Figure S1. Steady-state levels of HSP70-T636A and HSP70-T636D in HEK-293 cells.** Representative immunoblots of HSP70 steady-state levels with different amino acids at position 636 following the addition of cycloheximide (CHX) or MG132 with the percent change from time 0 at each 6 h time point. Cell extracts were separated via SDS-PAGE, and a Flag antibody detected the exogenous HSP70 levels.

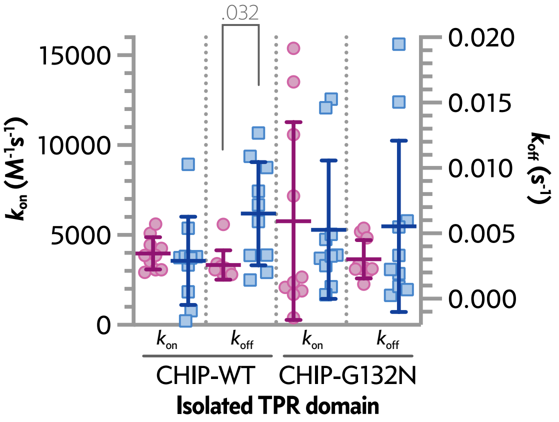


**Figure S2. G132N recovers increased $k_{off}$ between CHIP and pEEVD peptide.** Analysis of G132N using biolayer interferometry for wild type (WT) and G132N-CHIP-TPR measured by $k_{on}$ and $k_{off}$. Data represented by dot plot and summarized by the mean ± SD analyzed via two-way ANOVA: main effects of TRP construct, $F(1, 36) = 0.04545$, $P= 0.832$ and EEVD peptide, $F(1, 36) = 6.718$, $P = 0.014$, and an interaction effect between TPR construct and EEVD peptide, $F(1, 36) = 0.3208$, $P < 0.575$. TPR construct accounted for 0.11% of the total variation, while the EEVD peptide accounted for 16%. Post-test results of pairwise comparisons < 0.05 are included in the plot.

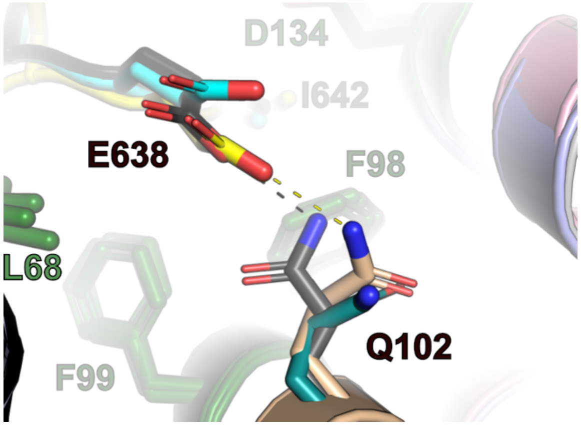


**Figure S3. Enhanced detail of CHIP Q102 interaction with HSP70 E638.** A close-up view of the differential interactions of HSP70 E638 and CHIP Q102. CHIP-TPR. Overlaid structures include CHIP-TPR in complex with the helical lid+tail domain of HSP70 (PDB ID 4KBQ: CHIP in light grey, HSP70 helical lid+tail domain in dark grey), in complex with EEVD (PDB ID 9DYA: CHIP in light blue, EEVD in yellow), and in complex with the pEEVD (9DYB: CHIP in light pink, pEEVD in cyan). Throughout, CHIP-TPR residues L68, F98, F99, and D134, which interact with EEVD, are colored forest green. The hydrogen bond O-N distance between the E638 side chain carboxylate and Q102 amide is 2.9 Å (black dashed line) for CHIP-TPR in complex with the helical lid+tail domain of HSP70 and 2.7 Å (yellow dashed line) for the CHIP-TPR in complex with EEVD. Movement of pEEVD and rotation of the CHIP-TPR Q102 side chain eliminates the hydrogen bond between the E638 side chain carboxylate and Q102 amide, with the O-N distance increasing to 5.8 Å.

### Supporting Tables

| Vector | Host | Experiment(s) | Tag | Key Mutation | Lab | AddgeneID |
|---|---|---|---|---|---|---|
| **CHIP-TPR** | Pro | Crystal Structures, Biolayer Interferometry | HIS | Wild-type | RCP | |
| **HSP70-lid/ tail** | Pro | Crystal Structures, Biolayer Interferometry | HIS | Wild-type | RCP | |
| **CHIP** | Pro | Fluorescent Polarization, In Vitro Ubiquitination | HIS | Wild-type | JCS | |
| **CHIP-K30A** | Pro | Fluorescent Polarization | HIS | K30A | JCS | |
| **CHIP- G132N** | Pro | Fluorescent Polarization, In Vitro Ubiquitination | HIS | G132N | JCS | |
| **CHIP TPR-CC** | Pro | Fluorescent Polarization, In Vitro Ubiquitination | HIS | UBOX deletion | JCS | |
| **CHIP TPR-UBOX** | Pro | Fluorescent Polarization, In Vitro Ubiquitination | HIS | CC deletion | JCS | |
| **CHIP CC-UBOX** | Pro | Fluorescent Polarization, In Vitro Ubiquitination | HIS | TPR deletion | JCS | |
| **CHIP TPR** | Pro | Fluorescent Polarization, In Vitro Ubiquitination | HIS | UBOX and CC deletion | JCS | |
| **WT HSP70** | Pro | In Vitro Ubiquitination | HIS | Wild-type | NGB | |
| **Empty Vector (pcDNA3)** | Euk | Co-Immunoprecipitation, Cell Proliferation | none | none | JCS | |
| **HSP70-WT** | Euk | Co-Immunoprecipitation, Cell Proliferation | FLAG | Wild-type | JCS | |
| **HSP70A** | Euk | Co-Immunoprecipitation, Cell Proliferation | FLAG | T636A | JCS | |
| **HSP70D** | Euk | Co-Immunoprecipitation, Cell Proliferation | FLAG | T636D | JCS | |
| **WT CHIP** | Euk | Co-Immunoprecipitation, Cell Proliferation | MYC | Wild-type | JCS | |
| **K30A CHIP** | Euk | Co-Immunoprecipitation, Cell Proliferation | MYC | K30A | JCS | |
| **G132N CHIP** | Euk | Co-Immunoprecipitation, Cell Proliferation | MYC | G132N | JCS | |
| **LgBiT-HSP70-WT** | Euk | NanoBiT | LgBiT Nanoluc | Wild-type | KMS | |
| **LgBiT-HSP70-A** | Euk | NanoBiT | LgBiT Nanoluc | T636A | JCS | |
| **LgBiT-WT HSP70-D** | Euk | NanoBiT | LgBiT Nanoluc | T636D | JCS | |
| **SmBiT-CHIP-WT** | Euk | NanoBiT | SmBiT Nanoluc | Wild-type | KMS | |
| **SmBiT-CHIP-K30A** | Euk | NanoBiT | SmBiT Nanoluc | K30A | JCS | |
| **SmBiT-CHIP-G132N** | Euk | NanoBiT | SmBiT Nanoluc | G132N | JCS | |

**Table S1: Listed are the vectors used in the study.** The experimental use, length, tag, mutation, and where the construct was acquired from are included for each construct. Vector sequences are available via Addgene. KMS = Kenneth Matt Scaglione, Duke University.

| | HsCHIP$_{21-154}$ + HsHsp70$_{634-641}$ | HsCHIP$_{21-154}$ + pT$_{636}$–HsHsp70$_{634-641}$ |
|---|---|---|
| **Data Collection** | | |
| Beam Line | ALS 4.2.2 | ALS 4.2.2 |
| Wavelength (Å) | 1.0000 | 1.0000 |
| Space group | $C222_1$ | $C222_1$ |
| **Cell dimensions** | | |
| *a*, *b*, *c* (Å) | 46.89, 74.36, 77.63 | 37.58, 45.93, 78.15 |
| *a*, *b*, *g* (°) | 90, 90, 90 | 90, 90, 90 |
| Resolution (Å)[a] | 39.67-1.89 (1.98-1.89) | 39.60-1.59 (1.64-1.59) |
| $R_{merge}$ [b] | 0.170 (1.014) | 0.092 (0.859) |
| $R_{meas}$ [c] | 0.198 (1.21) | 0.107 (1.013) |
| $CC_{1/2}$ | 0.988 (0.442) | 0.997 (0.612) |
| $<I/sI>$ | 7.05 (0.99) | 10.32 (1.25) |
| Wilson *B* factor (Å$^2$) | 14.66 | 13.58 |
| Completeness (%) | 98.93 (91.46) | 99.70 (97.38) |
| Redundancy | 3.8 (3.2) | 3.7 (3.5) |
| No. of reflections | 77,973 (7,819) | 127,365 (8,914) |
| No. of unique reflections | 20,723 (2,406) | 34,506 (2,570) |
| **Refinement** | | |
| No. reflections for refinement | 11,040 (1,242) | 18,531 (1,855) |
| $R_{work}/R_{free}$ | 0.189 / 0.234 | 0.173 / 0.213 |
| **Average *B* factors** | | |
| Protein | 18.18 | 16.53 |
| Water | 24.47 | 24.95 |
| Ions | 21.11 | 20.47 |
| **R.m.s deviations** | | |
| Bond lengths (Å) | 0.010 | 0.006 |
| Bond angles (°) | 1.160 | 0.780 |
| **Ramachandran plot statistics** | | |
| Favored regions % (#) | 99.2 (131/132) | 97.9 (137/140) |
| Allowed regions % (#) | 100.0 (132/132) | 100.0 (140/140) |
| Disallowed regions | 0.0 | 0.0 |
| **MolProbity validation statistics** | | |
| Cb deviations >0.25Å | 0 | 0 |
| MolProbity clash score | 1.85 | 1.80 |
| MolProbity clash percentile | 99$^{th}$ percentile (N=1,784, all resolutions) | 99$^{th}$ percentile (N=1,784, all resolutions) |
| MolProbity score | 0.95 | 1.00 |
| MolProbity score percentile | 100$^{th}$ percentile (N=27,675, all resolutions) | 100$^{th}$ percentile (N=27,675, all resolutions) |
| PDB ID | 9DYA | 9DYB |

**Table S2: X-ray data collection and structure refinement used to generate Figure 3.** [a] Values in parentheses are for the highest resolution shell. [b] The merging *R* factor measures the consistency between multiple measurements of the same reflection. [c] The corrected *R* factor was the final adjustment used for known systematic errors in the data.

| Name | Target | Type | Company | Reference Number |
| --- | --- | --- | --- | --- |
| DYKDDDDK Tag (D6W5B) | FLAG-Tagged HSP70 | Rabbit mAb (HRP conjugate) | Cell Signaling | 86861S |
| Myc-Tag (9B11) | MYC-Tagged CHIP | Mouse mAb (HRP conjugate) | Cell Signaling | 4040S |
| CHIP (C3B8) | CHIP | Rabbit mAb | Cell Signaling | 2080S |
| Hsp70/Hsp72 (C92F3A-5) | HSP70 | mAb | Enzo | ADI-SPA-810 |

**Table S3: Antibodies used in the study.**

| Figure | Type/Antibody | Image |
|---|---|---|
| 1I | IP: Stain-free | 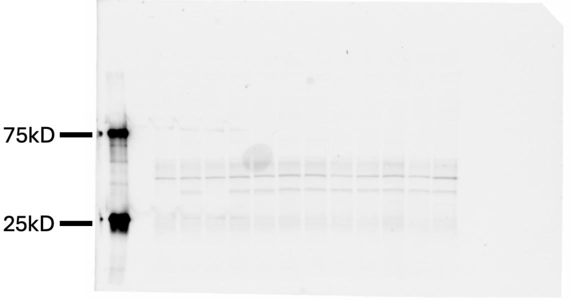  |
| | IP:Flag | 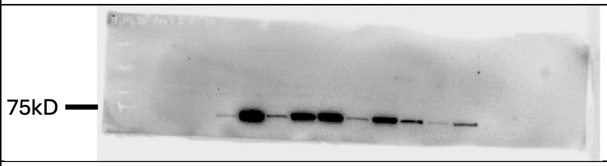  |
| | IP:MYC | 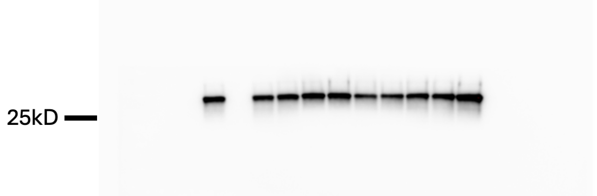  |
| | Input: Stain-free | 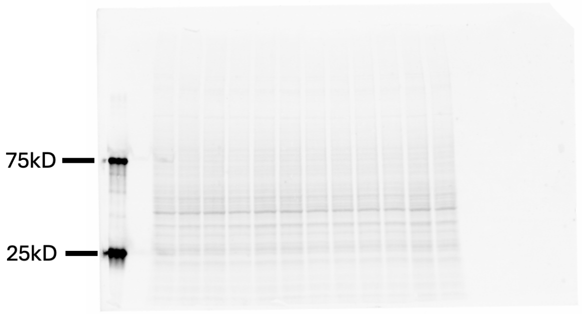  |
| | Input: Flag | 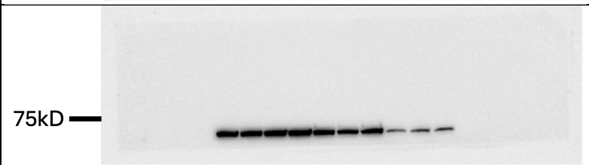  |
| | Input: MYC | 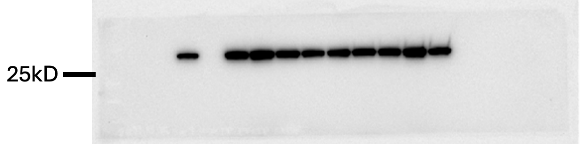  |

| | | |
|---|---|---|
| **2J** | Stain-free | 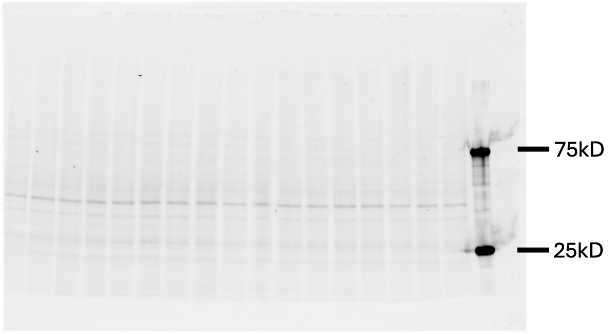  |
| | FLAG | 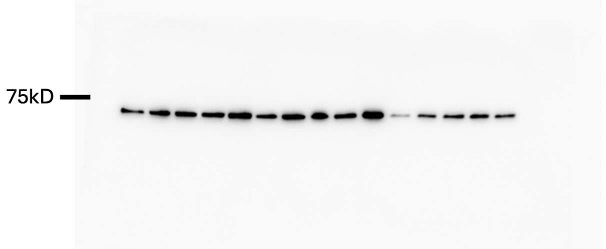  |
| **2K** | Stain-free | 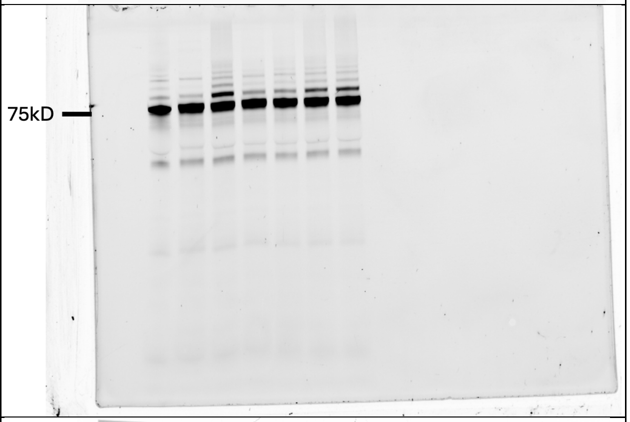  |
| **S1** | Stain-free | 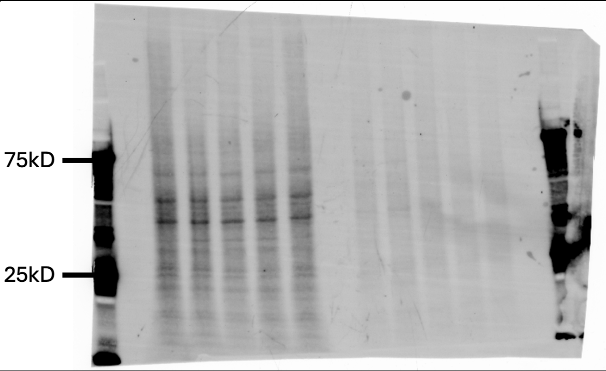  |

| | | |
|---|---|---|
| **S1A** (cont.) | Flag | 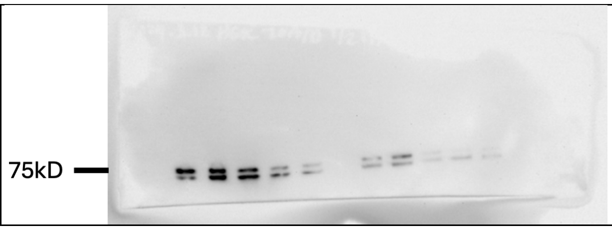 |

**Table S4. Stain-free and unmodified immunoblots.** The stain-free membrane before blocking and primary antibody and unmodified representative immunoblots are included for each figure that utilizes an immunoblot. The antibody used is included in the second column.
