## Supplementary material for "Phosphorylation-State Modulated Binding of HSP70: Structural Insights and Compensatory Protein Engineering": Table 1

### TABLES

| | HEK-293 | | COS-7 | |
|---|---|---|---|---|
| **Source of Variation** | **% of total variation** | **P value** | **% of total variation** | **P value** |
| CHIP | 0.07 | 0.997 | 0.12 | 0.993 |
| Time | 10 | 8.57E-03 | 31 | 2.27E-05 |
| HSP70 | 0.04 | 0.873 | 0.04 | 0.869 |
| CHIP x Time | 0.10 | 0.995 | 0.01 | 1.000 |
| CHIP x HSP70 | 0.02 | 1.000 | 0.09 | 0.996 |
| Time x HSP70 | 0.01 | 0.920 | 0.23 | 0.688 |
| CHIP x Time x HSP70 | 0.18 | 0.988 | 0.07 | 0.997 |

**Table 1. Three-way ANOVA on CHIP and HSP70 modifications and time with cell counts.** The table shows the percentage of total variation and P values for each source of variation, including single terms and interaction terms (denoted by 'x'), for both HEK-293 and COS-7 cell lines.
